# A Comparison of Deep Learning Architectures for Inferring Parameters of Diversification Models from Extant Phylogenies

**DOI:** 10.1101/2023.03.03.530992

**Authors:** Ismaël Lajaaiti, Sophia Lambert, Jakub Voznica, Hélène Morlon, Florian Hartig

## Abstract

To infer the processes that gave rise to past speciation and extinction rates across taxa, space and time, we often formulate hypotheses in the form of stochastic diversification models and estimate their parameters from extant phylogenies using Maximum Likelihood or Bayesian inference. Unfortunately, however, likelihoods can easily become intractable, limiting our ability to consider more complicated diversification processes. Recently, it has been proposed that deep learning (DL) could be used in this case as a likelihood-free inference technique. Here, we explore this idea in more detail, with a particular focus on understanding the ideal network architecture and data representation for using DL in phylogenetic inference. We evaluate the performance of different neural network architectures (DNN, CNN, RNN, GNN) and phylogeny representations (summary statistics, Lineage Through Time or LTT, phylogeny encoding and phylogeny graph) for inferring rates of the Constant Rate Birth-Death (CRBD) and the Binary State Speciation and Extinction (BISSE) models. We find that deep learning methods can reach similar or even higher accuracy than Maximum Likelihood Estimation, provided that network architectures and phylogeny representations are appropriately tuned to the respective model. For example, for the CRBD model we find that CNNs and RNNs fed with LTTs outperform other combinations of network architecture and phylogeny representation, presumably because the LTT is a sufficient and therefore less redundant statistic for homogenous BD models. For the more complex BiSSE model, however, it was necessary to feed the network with both topology and tip states information to reach acceptable performance. Overall, our results suggest that deep learning provides a promising alternative for phylogenetic inference, but that data representation and architecture have strong effects on the inferential performance.

## Introduction

Species richness varies greatly across taxonomic groups (G. E. Hutchison 1959), geological times (Barnosky et al. 2011) and geographical regions (Gaston and Blackburn 2000). It is generally accepted that these patterns in species richness emerge from variation in speciation and extinction rates in addition to dispersal and migration processes. For instance, important ecological phenomena such as the ‘Latitudinal Diversity Gradient’ (Hillebrand 2004) may be partly explained by variations in species net diversification rates defined as the balance of the speciation and the extinction rate (Mittelbach et al. 2007; Rolland et al. 2014; Pontarp et al. 2019). Our general understanding of the processes that cause these rates changes, however, is still very limited (Rabosky 2009a, 2009b; Condamine et al. 2013; Moen and Morlon 2014).

To better understand the drivers of variation in diversification rates and their consequences for biodiversity dynamics, it is crucial first to accurately estimate net diversification rates (Pyron and Burbrink 2013), and then decompose them into speciation and extinction rates (Stadler 2013). The latter is not trivial, as fossil data are rarely available and reconstructed phylogenies do not include information on extinct species. Indeed, without further constraints, the problem is ill-posed, meaning that different diversification dynamics can lead to the same extant phylogeny (Louca and Pennell 2020; Morlon et al. 2022). However, when making additional assumptions about the functional form of extinction and speciation rates over time, it is often possible to statistically estimate the parameters of these functions, and thus infer the path that led to the currently observed species (Nee et al. 1994; Etienne and Rosindell 2012; Morlon 2014).

Arguably one of the simplest diversification models is the Constant Rate Birth-Death (hereafter ‘CRBD’) model, which assumes that the speciation and extinction rates are both homogeneous across lineages and constant through time. Nee et al. (1994) showed that extinction and speciation rates in this model can be estimated based on reconstructed phylogenies with Maximum Likelihood using a distinct pattern called the “pull of the present”. In recent years, many studies have worked on extending this model to account for various types of rate heterogeneity as well as their potential drivers (Morlon et al. 2011; Etienne et al. 2012). These models commonly allow for variations of speciation and extinction rates through time (time-dependent models), across lineages (inhomogeneous models) as well as in dependence of environmental (Condamine et al. 2013) or biotic factors (Etienne et al. 2012). For instance, the time dependent Birth-Death model (Hallinan 2012) allows to consider homogenous changes (*e.g.* an exponential decay) of speciation and extinction rates over time (Rabosky and Lovette 2008; Morlon et al. 2011; Stadler 2011). Other diversification models allow lineage-specific shifts in diversification rates that can be either discrete, as can be expected to occur with the appearance of key innovations (Alfaro et al. 2009), or continuous, as can be expected to occur given the gradual evolution of phenotypes (*e.g.* the ‘ClaDS’ model; Maliet et al. 2019). The specific innovations or phenotypes that drive these shifts can be either implicit, as in the latter models, or explicitly modeled, as in State-dependent Speciation and Extinction (SSE) models. A particular example of this is the Binary State Speciation and Extinction model, hereafter ‘BiSSE’ (Maddison et al. 2007), which considers the effect of a binary state on speciation and extinction rates.

A constant challenge when developing new diversification models is establishing robust methods to fit them to data. For diversification models with low or moderate complexity, the likelihood (*i.e.* the probability to observe a reconstructed phylogeny given the model and its parameters) can be computed analytically or approximated numerically (see Nee et al. 1994 for the CRBD model and Maddison et al. 2007 for the BiSSE model). If the likelihood can be computed, model parameters can be inferred using either Maximum Likelihood Estimation (MLE; see for instance Ricklefs 2007) or Bayesian inference (*e.g.* Silvestro et al. 2011).

When the likelihood of a diversification model is analytically or numerically intractable, simulation-based methods such as Approximate Bayesian Computation (ABC) are typically being used. The ABC approach approximates the likelihood of a model by comparing model predictions to data via summary statistics (Csilléry et al. 2010; see Saulnier et al. 2017 for an example). Although ABC can successfully infer parameters from complex diversification models, it has two major drawbacks. First, ABC typically requires a large number of simulations, which is often computationally prohibitive in practical applications. Secondly, ABC suffers from the curse of dimensionality, meaning that the computational increases sharply with the dimensions of the summary statistics, which must be at least as large as the number of model parameters to ensure sufficiency (Beaumont 2010; Csilléry et al. 2010; Hartig et al. 2011). Together, both properties limit the applicability of ABC to complicated models with many parameters.

Recent advances in the field of deep learning algorithms provide an alternative for likelihood-free inference in phylogenetic inference. Unlike ABC, deep learning can easily deal with high-dimensional data and even benefit from it (LeCun et al. 2015), thus avoiding the necessity to find appropriate summary statistics. It was shown that deep learning approaches based on Convolutional Neural Networks (CNNs) can outperformed ABC for inference tasks in population genomics (Chan et al. 2018; Schrider and Kern 2018). Moreover, as pointed out by (Schrider and Kern 2018), neural networks are very flexible about their input data structure, possibly allowing a far greater or more diverse set of inputs than in traditional statistical models.

Unlike for ABC, however, there is little experience on using deep learning algorithms for parameter inference in diversification models, and there are a number of options to implement DL for phylogenetic inference. One of the first questions to solve is how to best encode phylogenetic data, which in turn affects the DL architectures that could be considered for training. Recently (Voznica et al. 2022) developed a full matrix representation for non-ultrametric phylogenies with the goal of fitting epidemiological models, and compared the performance of feed-forward neural networks (DNNs) and CNNs that were supplied either with this tree representation or with conventional summary statistics to likelihood-based inference techniques. (Lambert et al. 2022) adapted this approach with the goal of fitting birth-death diversification models to reconstructed (ultrametric) phylogenies, extended the matrix representation to the case of representing phylogenies with associated tip state data, and similarly compared the performance of DNN, CNN and MLE. These studies showed that the matrix encoding processed by CNNs performed very well, leading to gains in predictive performance compared to likelihood-based methods (Voznica et al. 2022). However, there are a number of further options regarding the encoding and the network architectures, and we conjectured that the optimal combinations of those would depend on the diversification model to be estimated.

When considering the choice of network architecture (Pichler and Hartig 2023), the simplest option would be a standard fully connected Deep Neural Network (DNN). The disadvantage of this option is that fully connected DNNs do not perform favorably with high-dimensional structured input data (such as phylogenies). Therefore, we can anticipate for successfully using DNNs, we would have to summarize the phylogeny’s shape (which by default is high dimensional with neighborhood relationships being represented as a graph) using a limited number of summary statistics (Saulnier et al. 2017), similar to the ABC approach. While possible, this might result in a loss of information for the inference, depending on whether those summary statistics are sufficient for the inference task.

A second option would be the use of Convolutional Neural Networks (CNNs). CNNs apply one or several filters that slide incrementally across the input data to detect spatial patterns. CNNs have shown great success for a variety of tasks, including for fitting birth-death models to pathogen or species phylogenies encoded in a full matrix representation (Voznica et al. 2022; Lambert et al. 2022). CNNs could also be used on reduced representations of the phylogeny, such as the Lineage Through Time (hereafter ‘LTT’) plot. This seems promising as it is known that the slope of the LTT is a sufficient statistics for homogeneous models (Nee et al. 1994; Ricklefs 2007).

When representing the phylogeny by its LTT, another option is the use of Recurrent Neural Networks (RNNs). The neurons of RNNs include a temporal feedback and, thanks to this feature, RNNs can process sequential data (*e.g.*, time series) more efficiently. A common and efficient RNN neuron architecture is the Long Short-Term Memory (LSTM) cell. RNNs based on LSTM cells can flexibly handle long and short term dependencies of the input data (Hochreiter and Schmidhuber 1997; Yu et al. 2019) and have been applied with success in many fields, including Natural Language Processing tasks (Huang et al. 2019), financial market forecasting (Bukhari et al. 2020) or phoneme classification (Graves et al. 2005).

Finally, when considering the nature of a phylogeny, Graph Neural Networks (GNNs) arise as a natural choice. GNNs generalize the idea of CNNs, which require Euclidean neighborhoods, to graphs (Scarselli et al. 2009; Zhang et al. 2019; Wu et al. 2021). GNNs are successfully used for many applications with a graph structure, *e.g.* predicting molecules properties (Gilmer et al. 2017; Mansimov et al. 2019). The potential advantage of using GNNs over CNNs is that GNNs can directly perform convolutions along the phylogeny’s topology, whereas the use of a CNN requires transforming the phylogeny into a Euclidean structure, which potentially distorts neighborhood relationships.

The hypotheses formulated in the previous paragraphs are based on the theoretical knowledge about DL architectures, but so far, no systematic comparison of the combination of phylogeny representation and neural network architecture has been performed to confirm these conjectures. For example, while CNNs feeding on encoded phylogenies were shown to infer rates with a good accuracy for different birth-death models (Voznica et al. 2022; Lambert et al. 2022), we might expect that this full encoding is redundant and thus suboptimal for homogeneous birth-death models, because for these models, all the relevant information is in the LTT (Lambert and Stadler 2013).

In this study, we systematically explore the interplay between model, data representation and network architecture when inferring parameters of diversification models using deep learning. As models, we considered the simple homogeneous CRBD model and the more complex inhomogeneous BiSSE model, with the idea that homogenous diversification models will require other data representations than inhomogeneous models. We represented phylogenies with either: LTTs, sets of summary statistics, phylogeny encodings from (Voznica et al. 2022 and Lambert et al. 2022), or phylogeny graphs. Then, these representations were combined with the suitable neural network architecture(s): summary statistics with DNNs, phylogeny graphs with GNNs, encoded phylogenies with CNNs, and LTTs with CNNs or RNNs. To evaluate the predictive performance of these deep learning inference methods, we compared the prediction errors of each method and broke them down into three terms: 1) variance, 2) uniform bias and 3) consistency bias (Smith and Rose 1995). As a reference, we compared the deep learning results to the MLE for these models, which is tractable in both cases.

Using this set-up, we ask three questions. First, can deep learning methods accurately infer rates from diversification models, as suggested by (Lambert et al. 2022), and if so, can these methods outperform MLE regarding the prediction error? We hypothesize that deep learning methods can theoretically outperform MLE as they can trade off bias against variance to minimize the total error. Secondly, what is the optimal representation of phylogeny data to infer rates for diversification models of different complexity? We expect, for example, that the LTT is a sufficient statistic for simple homogeneous models and is often preferable due to its simplicity. For more complex models and specifically for inhomogeneous models, on the other hand, we expect that simple representations such as the LTT will fail to provide the neural network with the necessary information. More complex representations are necessary for optimal inferential performance. Lastly, we ask how the choice of the neural network architecture in combination with the chosen phylogeny representation affects the inference. We hypothesize that the more complex the phylogeny representation, the more important the neural network architecture becomes.

## Methods

### Diversification Models

We chose two established diversification models as case studies to test the performance of deep learning algorithms for phylogenetic inference: 1) the relatively simple and homogeneous CRBD model, and 2) the more complex inhomogeneous BiSSE model. The rationale for choosing these two models was to have two models with tractable likelihoods but different complexity, and especially different in their (in)homogeneity, as we conjectured that homogenous BD will profit less from detailed representations of the phylogeny, given that it is known that all information for their inference is contained in the LTT.

The Constant Rate Birth-Death (CRBD) model is one of the simplest conceivable diversification models. It has two parameters, the speciation rate (λ) and the extinction rate (μ), that are homogenously constant over time. The likelihood of this model is well-known (Nee et al., 1994) and depends only on the phylogeny’s branching times and not on its topology. To estimate diversification rates with the MLE, e used the APE R package (Paradis et al. 2004).

In the Binary State Speciation and Extinction (BiSSE) model, lineages can alternate between two states (0 and 1), where each state has its own speciation and extinction rate. The switch through time between the two states translates character transitions (*e.g.* sexual to asexual reproduction) that can have an impact on diversification rates. The model has 6 parameters: 2 speciation rates (λ_0_, λ_1_), 2 extinction rates (μ_0_, μ_1_) and 2 two transition rates (q_01_ for transition 0 → 1; q_10_ for transition 1 →0). As for the CRBD model, the likelihood of the BiSSE model can be computed (see Maddison et al., 2007). We imposed four constraints on the model to simplify inference, thus decreasing the number of free parameters from six to two. The constraints are: 1) λ_1_ = 2λ_0_; 2) μ_0_ = 0; 3) μ_1_ = 0; 4) q_01_ = q_10_. Constraint 1) ensures that states 0 and 1 have different speciation rates, 2) and 3) make the model pure birth, and 4) is an assumption of symmetry that makes the probabilities to switch from one state to another equal. Lambert et al. (2022) also used the CRBD model, and a less constrained version of BiSSE. Here we simplified the model to focus on the comparison of network architectures and data representation. To estimate diversification rates with the MLE, we used the Diversitree R package (FitzJohn 2012) and constrained the likelihood with the 4 constraints listed above.

To train the neural networks, we simulated 100,000 phylogenies for the CRBD model and 1,000,000 phylogenies for the BiSSE model using the Diversitree library (FitzJohn 2012) in R. We assumed complete sampling of the phylogenies. The latter assumption (no missing species) could be relaxed (Lambert et al. 2022), but here our focus is on the comparison between inference methods. For the CRBD model, we draw the underlying parameters as follows: 1) we draw uniformly λ in [0.1,1.0]; 2) we draw uniformly the turnover rate ε in [0, 0.9] from which we compute the extinction rate μ = ελ ∊ [0, 0.9λ]. By doing so, we avoid the critical case where λ ≲ μ (*i.e.,* speciation rate is inferior to or equivalent to the extinction rate). For the BISSE model, we took λ_0_ ∊ [0.1,1.0] to stay in the same range as for the CRBD model and q_01_ ∊ [0.01,0.1] to ensure that one state is not overly represented compared to the other one.

### Phylogeny Representation

Most of the considered deep learning architectures cannot process phylogenies as inputs, but require reformatting the phylogenies into a more regular data structure. Here, we explain all options that we considered (see also Fig. 1).

**Figure 1.**
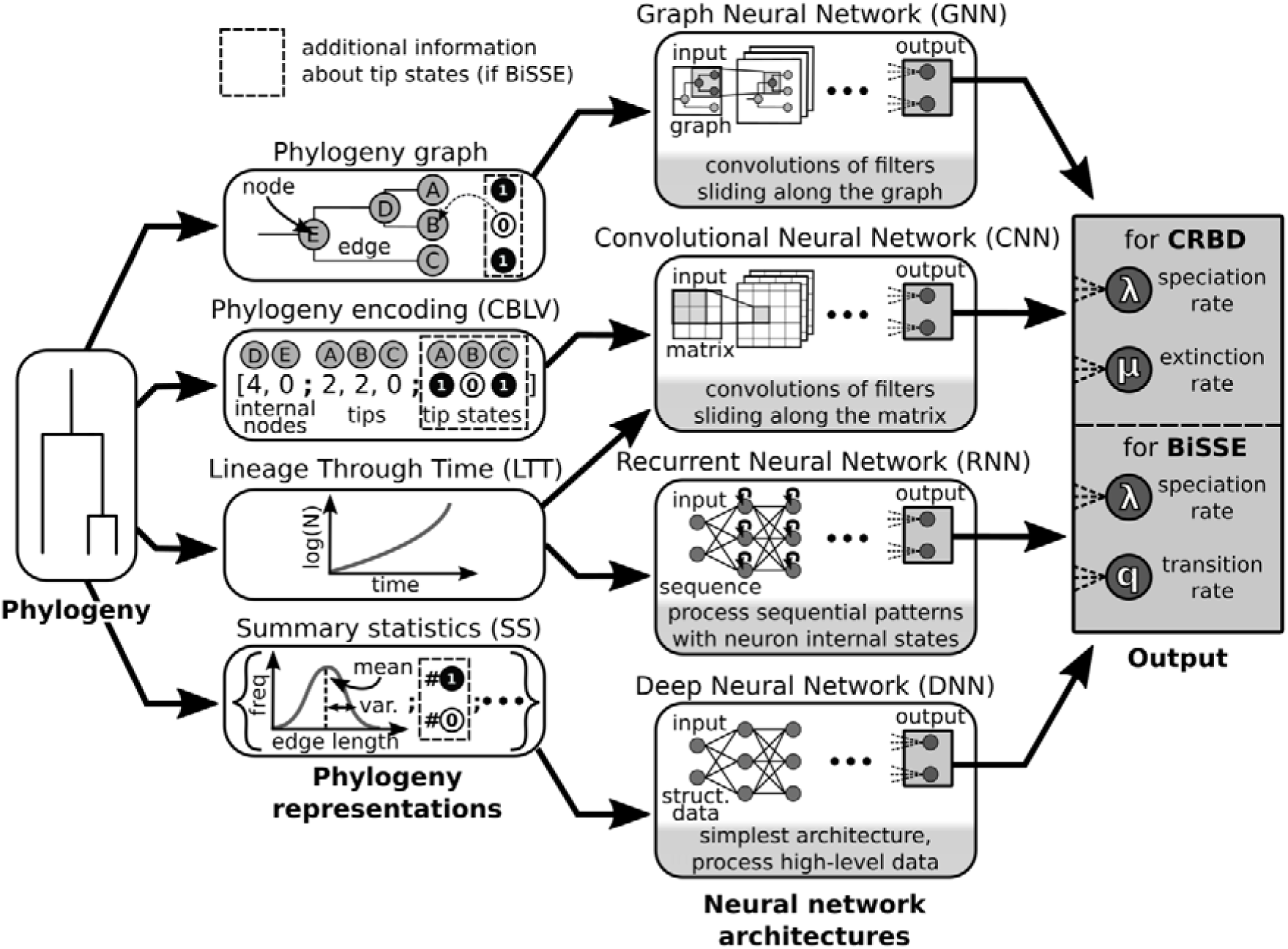
Combining phylogeny representations with neural network architectures. The five different combinations considered between phylogeny representations and neural network architectures are indicated by arrows from the first column to the second one. The rationales behind these five combinations are the following: 1) for phylogeny graph with GNN, GNN is the only architecture able to process graphs; 2) for CBLV with CNN, CNNs are able to detect patterns that are expected in the phylogeny encoding (Voznica et al. 2021); 3) for LTT with CNN, CNNs can detect patterns in the slope of the LTT; 4) for LTT with RNN, RNNs are designed to process time series; 5) SS with DNN, SS do not have spatial or temporal hierarchical order and therefore can be processed by a simple architecture. The third column describes the output of the inference methods for the two diversification models examined (CRBD and BiSSE).

#### Summary statistics

Arguably the most basic option is to represent the phylogeny by a number of summary statistics. We used a set of 84 summary statistics inspired from the set of (Saulnier et al. 2017). Those summary statistics can be split into three groups:

1. 8 statistics related to phylogeny topology (*e.g.* ratio of the width over the depth of the phylogeny);
2. 25 branch lengths statistics (*e.g.* median of all branch lengths);
3. 51 LTT statistics (binned LTT coordinates and LTT slopes).

The list of the summary statistics and the changes compared to (Saulnier et al. 2017) are detailed in the Table S1 of the Appendix.

For the BiSSE model we use the ratio of the number of tips in state 1 over the total number of tips as an additional summary statistic.

#### Lineage Through Time (LTT)

LTTs illustrate the increase in lineages over time. For a phylogeny of n tips, the LTT is composed of n-1 points where each point is defined by two coordinates: time (t, abscissa) and the number of lineages (N, ordinate). For a binary tree where each branching event results in two daughter lineages, the LTT can be compressed to a 1-d array without loss of information, by throwing away the number of lineages and keeping only the times of speciation events.

#### Phylogeny encoding (CBLV)

Yet another alternative is to encode the phylogeny in a real-values vector of length 2n-1 named the ‘Compact Bijective Ladderized Vector’ (‘CBLV’, see Voznica et al., 2022). Each value of the vector corresponds to one node (internal node or tip), thus there are n values for the n tips and n-1 values for the internal nodes n-1, which result in a vector of 2n-1 values. The encoding is done in two steps: 1) phylogeny ladderization, and 2) phylogeny traversal. The principle of ladderization is to order each node’s children, such that the encoding is bijective (*i.e.*, one CBLV maps exactly to one phylogeny and reversely). Here we ladderized the phylogeny such that for each node the left child is the child which is further from the root. On the ladderized phylogeny, we perform an inorder traversal using a classical recursive algorithm. If the visited node is an internal node or the first tip is visited, its distance to the root is added to the vector. Otherwise, its distance from its parent node is added to the vector. Moreover, for the BiSSE model we add the n tip states to the vector. The tip states are ordered according to the phylogeny traversal. Note that in the original method from Voznica et al. (2022), the tip state information was added by adding one row to the distance vector, thus resulting in a two-row matrix (first row for distances, second row for tip states), while here we concatenate two 1-d vectors resulting in a 1-d vector. We found that this modification did not affect the results.

#### Phylogeny as a graph

The main motivation for the previous CBLV encoding is to transform the phylogeny into a regular Euclidean format that can be easily processed by CNNs. The disadvantage, however, is that neighborhood relationships of the graph can be distorted in this process. Recent research in the field of machine learning suggests that in such a case, it is often better to train neural networks directly with the original graphs. This is done by so-called Graph Neural Networks or GNNs, which extend the CNN idea to graphs. Here, we use the Pytorch Geometric framework (Fey and Lenssen 2019) and the GraphNeuralNetworks Julia package base on the Flux framework (Innes 2018) to represent the phylogeny as graph and train GNNs. We provide the GNN with the phylogeny’s topology and 4 attributes per node: distance to the root and the lengths of the 3 edges linked to the node (1 incoming edge: parent → node; 2 outcoming edges: node →child_1_ and node → child_2_). Moreover, for the BiSSE model we add as an attribute the tip states (0 or 1 for tips, -1 for nodes whose state is unknown).

### Neural Network Architectures

For predicting model parameters from the formatted phylogenies, we considered four different neural network architectures: DNN, CNN RNN and GNN (see also Fig. 1). All architectures were built within the torch framework (Falbel and Luraschi 2019) in R, except GNNs which were built with Torch in Python because the PyTorch Geometric library (Fey and Lenssen, 2019) was needed for GNNs. Hyperparameter tuning for each architecture was performed by hand (see Appendix Tables S0-S5).

### Training Neural Networks

To train and validate the performance of the networks, we simulated 100,000 phylogenies under the CRBD model and 1,000,000 phylogenies under the BiSSE model. For both models phylogenies were split randomly in three groups: 1) a training set to train the neural networks (90% of the phylogenies); 2) a validation set used to quantify performance of the neural network during the training (5% of the phylogenies) and 3) a test set used to quantify the performance of the neural network after the training (5% of the phylogenies). These split sets were the same for all neural network architectures, so all of the architectures are trained and tested with the same phylogenies. Moreover, the Maximum Likelihood Estimation (‘MLE’) was tested on the phylogenies of the test set to fairly compare neural networks and MLE.

Training of the networks followed the standard practice of dividing the training data into small batches (typically 64 phylogenies per batch, for more details see the Appendix). During optimization, batches are successively provided to the network and its weights are adjusted to reduce the prediction error (defined as the mean squared error). Once the network has been fed with the whole training set, the weights are frozen and the performance of the network is evaluated on the validation set. This process of training and validation is called an ‘epoch’. We repeated this process until the prediction error on the validation set stopped to improve (early stopping). This ensures that the network does not overfit the training data, which tends to improve the performance of the model on new data (Bengio 2012). In addition to early stopping, we also set at each new epoch a small proportion of the network weights to zero (dropout, set to 1%), another standard method to avoid overfitting. After the training of the networks, we evaluated their performances on the test set.

### Interpreting Predictions Error

Once the neural networks were trained, we use them to infer the parameter values of the models from the test data (the phylogenies in the hold-out). We also calculated the Maximum Likelihood Estimate for these phylogenies to establish a baseline. Thus, for each method (the neural networks or the MLE) we have the predicted model parameters (*y_pred_*) *vs.* the true values (*y_true_*).

To better understand the origin of deviations between the true and predicted parameter values, we divide the prediction error into stochastic error and bias according to Theil coefficients (Smith and Rose 1995). The idea of this approach is that the sum squared differences 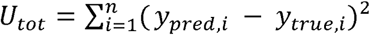 can be expressed as a sum *U_tot_ = U_uniformBias_ + U_consistencyBias_ + U_variance_* where: 1) 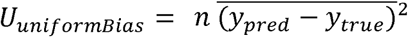 corresponds the systematic error; 2) 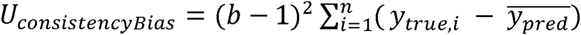 is the slope of the linear regression of *y_pred_ vs. y_true_* and 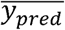 the mean of the predicted values; 3) 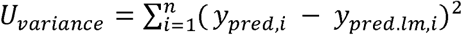 is the stochastic error, measured by the variance of the *y_pred_* around the regression slope *y_pred.lm,i_*. Lastly, we normalize each term by the sum squared of the true parameter values: 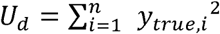. The error decomposition is illustrated in Figure 2.

**Figure 2.**
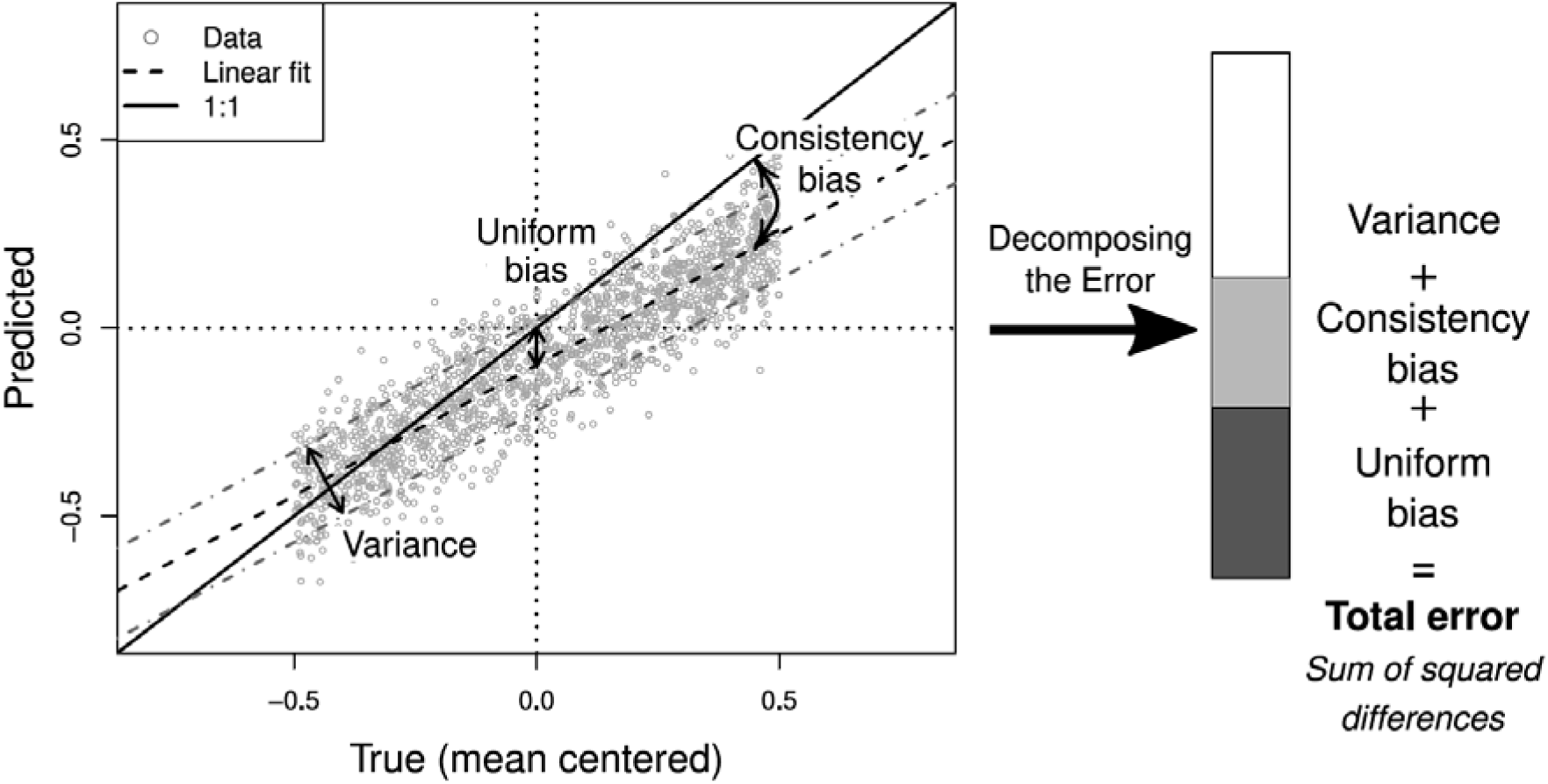
Decomposition of the prediction error. The total error defined as the sum of squared differences is split in three terms also known as the Theil’s coefficients: 1) variance that describe the width of the points distribution across the linear fit, 2) the uniform bias which quantifies the systematic error and 3) the consistency bias which expresses how far the slope of the linear fit is from 1 (Smith and Rose 1995). For the example we generate synthetic data such that: Predicted ∼ 0.7·True + N(μ=0.05,σ^2^=0.1) with 1,500 True values uniformly distributed in [0,1].This decomposition of the prediction error is summarized by a stack barplot (left, not to scale; for another example see Fig. 3). Note that the True axis has been mean centered such that the uniform bias can be measured at x=0.

To fairly compare neural networks and the Maximum Likelihood Estimator, we restricted the parameter space to test the inference models to an inner domain of the explored parameter space. Indeed, during the training phase the neural networks learn to not predict values outside the parameter space they have seen, whereas the Maximum Likelihood cannot do so. Thus, when estimating parameters close to the boundaries of the explored parameter space, the variance of the neural network predictions artificially decreases due to the specific set up of the phylogeny simulations.

## Results

Overall, we find that deep learning methods show good performances as they predict rates with an error comparable to MLE in most cases (Fig. 3). Some deep learning methods even outperformed MLE (CNN and RNN with LTTs for the CRBD model, see Fig. 3a,b). From theory, one would expect that lower total error of DL methods can be explained by the bias-variance trade-off, which allows DL methods to trade off bias against a lower total error (*e.g.* Pichler & Hartig, 2023). Looking at the decomposition of the error, however, we do not find that the deep-learning methods show greater bias than the MLE (*e.g.* DNN with SS vs. MLE for the CRBD model in Fig. 3a,b). The only notable exception to the good performance of the deep learning models was the GNN as well as the RNN and CNNs fed with the LTT for the BISSE model, which all performed notably poorer than the MLE.

**Figure 3.**
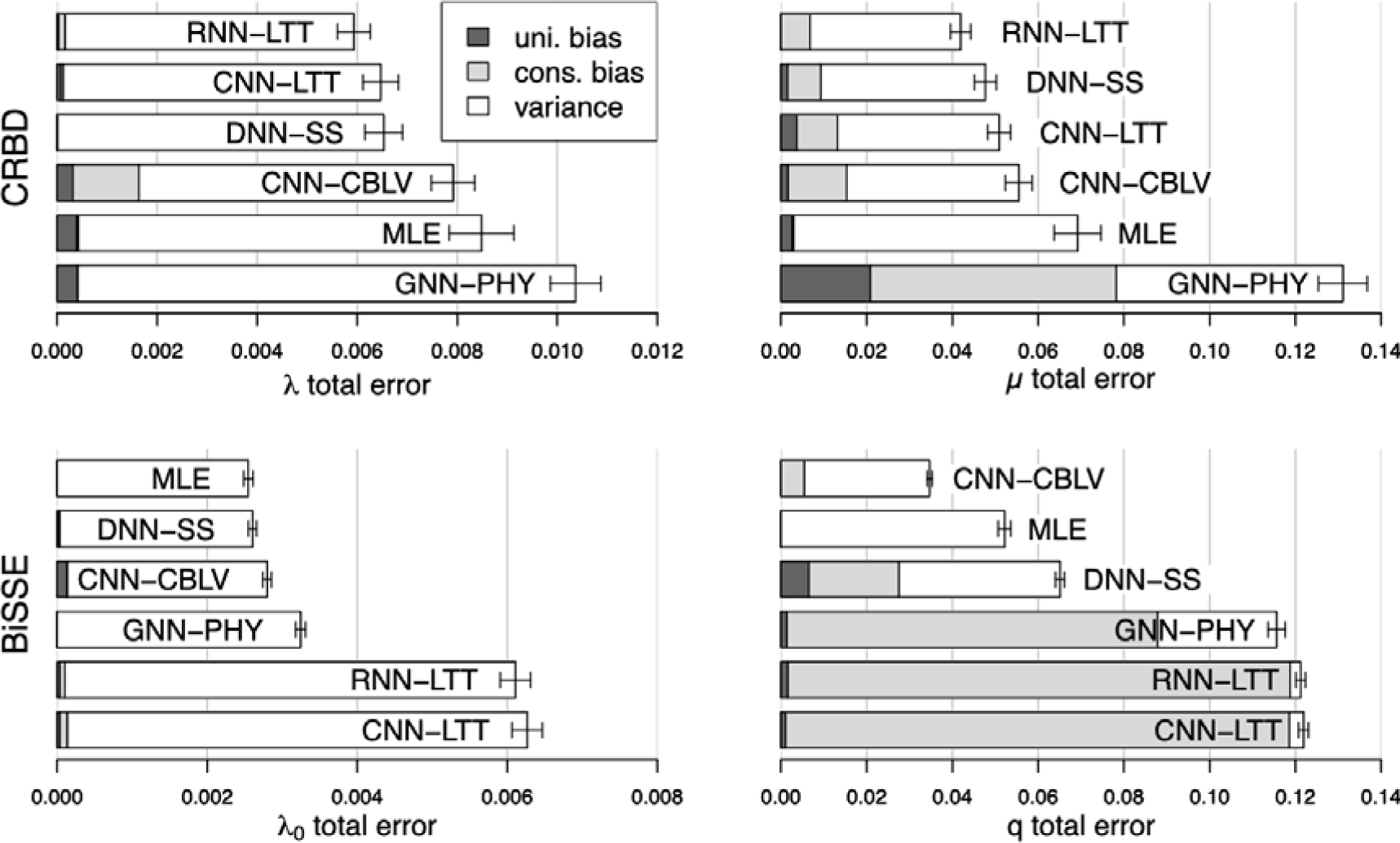
Prediction error of deep learning methods and MLE for the CRBD (a-b) and the BiSSE (c-d) model. From the CRBD we infer: a) the speciation rate (λ) and, b) the extinction rate (μ). From the BiSSE model we infer: c) the speciation rate of state 0 (λ_0_) and, d) the transition rate (q), the other parameters are constrained (see Methods for more details). The deep learning methods evaluated are the following: DNN with summary statistics (DNN-SS), CNN with LTTs (CNN-LTT), RNN with LTTs (RNN-LTT), CNN with encoded phylogeny (CNN-CBLV) and GNN with phylogeny graphs (GNN-PHY). The dataset contains 100,000 phylogenies for the CRBD model and 1,000,000 for the BiSSE model. The total summed-squared error is split in three terms: variance, uniform bias and consistency bias (for more details see Fig. 2 or Smith and Rose 1995). The error bars correspond to the 95% confidence interval.

Our second question was to understand how the optimal phylogeny representation depends on the complexity of the underlying diversification model. Our findings show that for the CRBD model, the best models are based on the LTT (CNN-LTT and RNN-LTT). Those models can even outperform the MLE (Fig. 3a,b). In contrast, inference methods relying on more complex phylogeny representations have a higher prediction error. This is especially clear for the two methods based on the most complex representations: CNN with full phylogeny encodings and GNN with phylogeny graphs. In sum, the more redundant information is added in the data, the worse the deep learning methods perform.

For the BiSSE model, we found a different behavior. CNN with encoded phylogenies and GNN with phylogeny graphs outperform simpler LTT-based models (Fig. 3). To understand which information is missing in the LTT and thus is responsible for the increased performance of the CNN with encoded phylogenies and GNN with phylogeny graphs compared to LTT-based models, we provided neural networks with data of more fine-grained increasing complexity. Specifically, we compared a CNN fed with LTTs as a baseline (1) with a CNN and three increasingly detailed encoded phylogenies (2-4). In the latter, we varied 2 features: the number of tip states and the location of tip states in the phylogeny. Thus, the 3 options are: excluding all tip states information (2); including information about the tip states, but not their location (3); including information about the tip states and their location (4; see Fig. 4c). To sum up, we compare 4 options: (1) no information about phylogeny’s topology, tip state numbers and location, (2) information about topology but not about tip states, (3) information about phylogeny topology, tip state numbers but not about their location, and (4) information about phylogeny topology, tip state numbers and their location.

**Figure 4.**
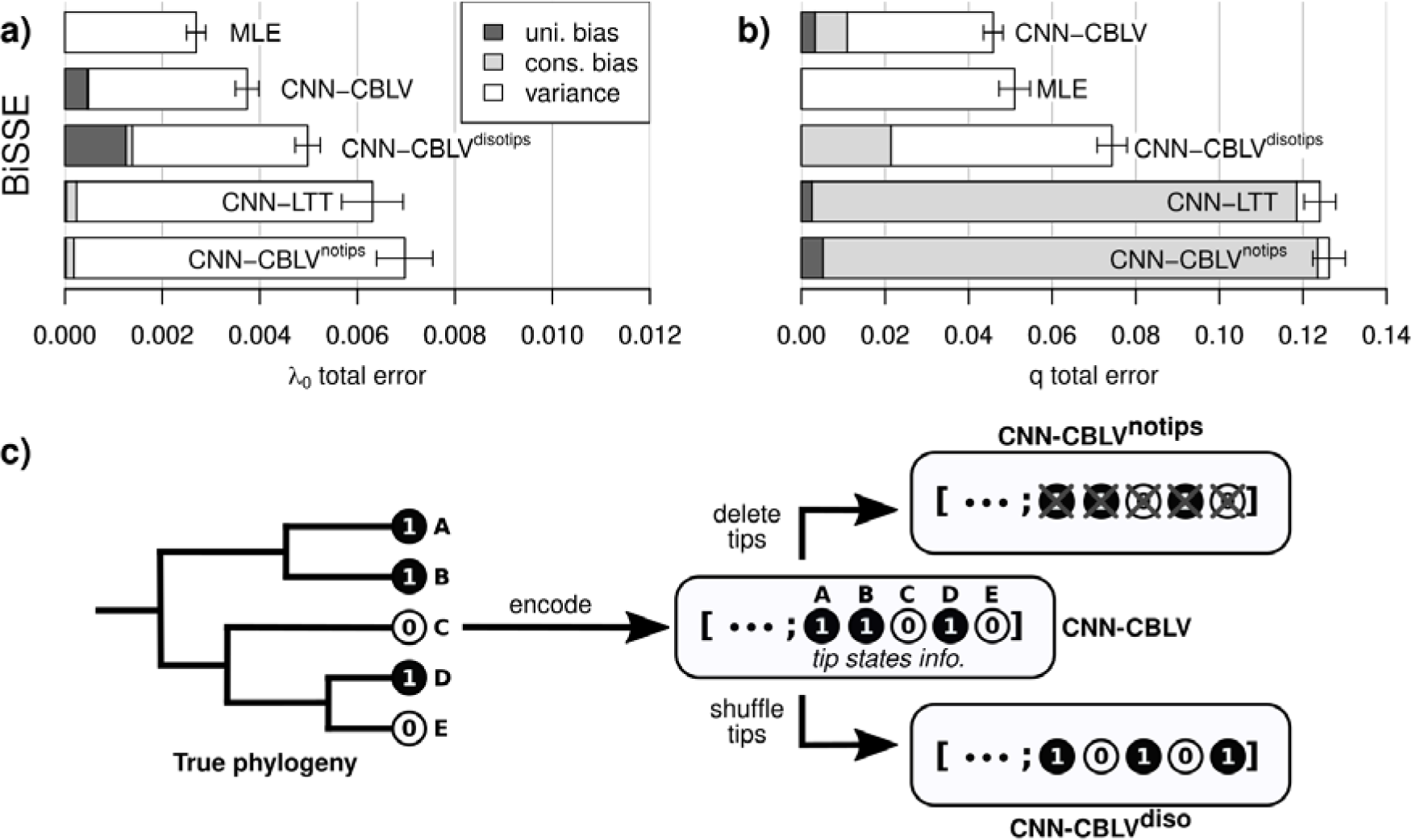
Prediction error of deep learning methods using different levels of tip states information for the BiSSE model. We infer: a) the speciation rate of state 0 (λ_0_) and, b) the transition rate (q), the other parameters are constrained (see Methods for more details). c) The deep-learning methods evaluated are CNN with encoded phylogenies: excluding tip states information (CNN-CLBV^notips^), containing tip states but randomly shuffled (CNN-CBLV^disotips^), and containing tip states in order (CNN-CBLV) and are presented in panel c). Additionally, we consider CNN with LTT to have a reference of another deep learning method that does not use tip states information, and MLE which is our baseline. The dataset for training and testing neural networks contains 100,000 phylogenies. The total summed-squared error is split in three terms: variance, uniform bias and consistency bias (for more details see Fig. 2). The error bars correspond to the 95% confidence interval.

This analysis confirms that the LTT alone is insufficient to infer rates accurately, as can be seen by the CNN and RNN combined with LTTs being among the deep-learning methods with the highest error (Fig. 3c,d). Moreover, the predictive performance increases with each information that was added about tip states, suggesting that each of the intermediate steps considered loses information (Fig. 4).

When looking at the influence of the neural network architecture, we see that this choice interacts with the data representation: for simple data representations of the phylogeny, CNN and RNN combined with LTTs achieved a similar prediction accuracy in all cases (Fig. 3), and they were overall the best option for the CRBD model, whereas for the more complicated BiSSE model, the even simpler DNN architecture with summary statistics outperformed the LTT-based architectures, whereas the CNN with encoded phylogenies performed best overall, including the MLE (Figs. 3, 4). The information provided to this architecture is equivalent, although differently structured, than the information provided to the GNN. The latter, however, performed notably poorer.

## Discussion

The goal of this study was to compare the ability of different deep learning architectures to infer parameters of diversification models from reconstructed phylogenies. Our main findings are that deep learning methods are surprisingly accurate for this task and can even outperform the MLE in some cases. Looking in more detail at the deep learning method, we find that the optimal phylogeny representation and network architecture depend on the diversification model. For the simple homogeneous CRBD model, we found that feeding the networks with the LTT was the main factor that improved inference, presumably because the LTT is a sufficient statistic and thus provides less redundant information to the networks. For the inhomogeneous BISSE model, it was optimal to provide the data in its most complex form, and the choice of the network architecture played a far greater role. We speculate that these results will generalize: if simple sufficient statistics exist for a model, these should be used and the choice of the network architecture is likely secondary. If no simple sufficient data representation exists for a given model, more complicated network architectures have to be used, and the choice of the network architecture becomes more critical.

Note that, in line with these thoughts, the performance of the DNN with SS for the BISSE model might be improved by adding more informative summary statistics that encode tip state information. This idea is supported by our observation that predictions of CNNs with CBLV outperform predictions of DNN with SS for the transition rate by a great margin, suggesting that tip state information that are contained in the CBLV encoding is critical for the parameter inference.

### Total Error and Bias

We ranked the different inference methods primarily by their total error, which sums up bias and variance (Smith and Rose 1995). This is in line with general practice in machine learning. To achieve a lower error, machine learning methods often trade off bias against variance (Pichler and Hartig 2023), whereas in statistics, bias is often seen as more crucial than variance, because an unbiased estimator allows a field to accumulate evidence over time (Shmueli 2010). Yet, given that there are usually no independent replicates of a phylogeny, we find it defensible to use the total error as the primary performance metric. Moreover, as we discuss below, DL algorithms did not generally exhibit larger bias than the MLE, suggesting that the problem of bias may be less severe than one might have anticipated.

Our results regarding total error on inferred parameters support earlier findings of (Voznica et al. 2022), who also reported that deep learning methods can outperform state-of-the-art methods when inferring model parameters from phylogenies (BEAST 2 in his study, MLE in our study). Contrary to the general idea of the bias-variance trade-off expectations, we did not find that deep learning methods generate this performance advantage over the MLE by leveraging the bias-variance trade-off. Indeed, several deep learning methods often had a lower total error and a lower bias than MLE (where we expected that they traded off variance against a greater bias to achieve lower error). To explain these findings, we speculate that the MLE apparently also has some small-sample bias for the CRBD model, especially for low net diversification rates (i.e. μ ≾ λ).

### Explaining the poor performance of the GNNs

Against our expectations, GNNs did not outperform other DL architectures and specifically the CNN with encoded phylogenies. This is surprising given the generally positive results in the machine learning literature about GNNs (Zhang et al. 2019), and the fact that GNNs are fed with the most accurate and natural representation of the data, which is the phylogeny itself. We speculate that our results can be explained by two well-known limitations of GNNs: ‘hop neighborhood’ (Nikolentzos et al. 2020) and ‘over-smoothing’ (Oono and Suzuki 2019).

First, ‘hop neighborhood’ refers to the fact that, in a GNN, at each new layer, node attributes are updated by aggregating their neighbor attributes (Scarselli et al. 2009). Thus, after k layers, each node aggregates only nodes that can be reached in k or less ‘hops’. Thus, node attributes only contain information about local graph structures but do not encode macroscopic graph information (Kriege et al. 2018). We speculate that for inhomogeneous birth-death models, the most important info about parameters related to the inhomogeneity is contained in subclades that have diversified long ago, and thus typically have large graph distances. In such cases, the “short-sightedness” of GNNs might be a major limitation.

We tested to increase the number of GNN convolutional layers from 2 to 5 (for the exact architecture see Appendix Table S5-6) but it did not lead to a performance improvement. Indeed, the naïve way to counteract this limitation is to increase the number of GNN layers as the size of the ‘k-hop neighborhood’ increases with k the number of layers. However, by doing so one will encounter the second limitation: ‘over-smoothing’ (Oono and Suzuki 2019): when stacking many layers in a GNN, node attributes become indistinguishable from one another due to the cumulative aggregations occurring at each layer. This phenomenon is referred to as ‘over-smoothing’ (Li et al. 2018). Thus, adding layers in GNNs often results in a deterioration of predictive performances (Li et al., 2018).

The combination of the two latter limitations might explain why ‘classical’ GNNs failed to infer rates accurately from reconstructed phylogenies. We speculate that our results regarding GNNs could be improved by considering specific GNN variants designed to overcome these limitations (for over-smoothing see Li et al. 2019; and for hop neighborhood see Nikolentzos et al. 2020), and encourage further research in this direction.

### Designing Efficient Deep-Learning Methods for Rate Inference

Our results reinforce previous analyses (Voznica et al. 2022; Lambert et al. 2022) suggesting that deep learning methods are a viable alternative to infer birth-death rates from phylogenetic data. This is particularly interesting for cases where likelihoods are not tractable or hard to compute. We refine these insights by demonstrating that the performance of the deep learning methods may strongly depend on data representation and network architecture. Regarding the complexity of the phylogeny representation, we find that avoiding to provide redundant data seems to help the algorithms. For instance, for homogeneous models, the topology of the phylogeny does not influence the likelihood (Lambert and Stadler 2013). Thus, the LTT provides a less redundant representation of the phylogeny while still containing all information useful for the inference (sufficient statistic). For more complex diversification models, in particular for models with intractable likelihoods, it will likely be harder to find appropriate sufficient statistics, and deep learning methods that use the whole phylogeny might be needed. For the BiSSE model, we found that the fully encoded phylogeny (Voznica et al. 2022) containing phylogeny topology and tip states information was the best choice among the representations that we considered. In short, choosing a good representation of the data is important: over-complicated representation either costs unnecessary computational power or deteriorates the accuracy of the predictions for simple diversification models, while choosing an over-simplified representation for a complex diversification model may lead to poor predictions as the representation does not contain enough information to predict parameters properly.

Once a suitable phylogeny representation has been found, the appropriate neural network architecture has to be selected. First, the choice of the architecture is restricted by the data type *e.g.*, a time series can go either with a CNN or an RNN but do not fit with a DNN which is not designed to detect patterns or to process temporal data. Secondly, the importance of the choice of the network architecture among the remaining possibilities is dependent on the complexity of the phylogeny representation: the more complex the representation, the more important the choice of the architecture is. In other words, when data become more difficult to process, more attention should be paid to the choice of the neural network architecture that will process the data. In theory, network architectures could be further fine-tuned regarding parameters such as the width and depth of the network layers, activation functions, training parameters and regularization settings. Due to computational reasons, we did not perform such a hyperparameter tuning, but rather set the different classes of network architectures to sensible default settings.

### Outlook and Further Research

While our results are encouraging regarding the use of deep learning models for parameter inference, they are at this stage only a proof of concept, because both diversification models considered here are still relatively simple. Moreover, to keep the analysis and training times tractable, we simplified the inhomogeneous BiSSE model by assuming four constraints on the rate parameters. We hold that these choices were appropriate for the purpose of this study, which was to establish general principles of the problem, but it is clear that the practical benefit of this methods would be to perform inference on more complicated diversification models with intractable likelihoods and probably a higher number of parameters (such as, *e.g.*, Hagen et al. 2021). How well the deep learning methods perform in this scenario remains to be explored by further research. Results in (Voznica et al. 2022) and Lambert et al. (2022) suggest that the deep learning methods still perform well when increasing model complexity.

Moreover, inferring parameters with deep learning methods has a number of drawbacks compared to traditional MLE approaches. Most importantly, training networks requires generating a large set of simulated phylogenies under the considered diversification model. To avoid that each user has to undergo this process, it seems obvious that already trained neural networks must be provided, but those would be likely still much larger and possibly less portable than current statistical procedures.

## Conclusions

In conclusion, our study supports the idea that deep learning methods can be used to infer diversification rates from reconstructed phylogenies. In our results, they often showed similar and sometimes even better performance than the MLE. Thus, deep learning methods offer a promising approach to likelihood-based statistical inference, in particular for models whose likelihoods are intractable or hard to compute. However, our results also provide a warning that deep learning methods are very diverse and the choice of the method (*i.e.* the combination of a phylogeny representation and a neural network architecture) must be adjusted to the model considered. If that can be achieved, however, they may be instrumental in expanding our options to perform statistical inference for complicated macroevoluationary models.

## Supplementary Materials

The code to reproduce the results in this study is available at https://github.com/ilajaait/phylo-inference-ml. Simulations data and the online appendix are available at https://doi.org/10.5061/dryad.7h44j0zzq.

## Acknowledgements

We thank Maximilian Pichler and Sonia Kéfi for helpful discussions and comments on the manuscript. The design of our study profited from the results of a MSc Thesis by Felix Gottschlich at the University of Regensburg that examined the possibility to use GNNs for phylogenetic inference.

