## Supplementary material for "A Comparison of Deep Learning Architectures for Inferring Parameters of Diversification Models from Extant Phylogenies": Online appendix

### *List of summary statistics*

The changes in summary statistics compared to for (Saulnier et al., 2017) occur in the summary statistics of the LTT plot. Here, in addition to 40 statistics to bin the LTT in 20 points, we added 10 slopes of the LTT instead of 2 slopes for (Saulnier et al., 2017). To compute the 10 slopes, the LTT is first cut into 10 equal parts regarding time; then points of each part are fitted in scale independently with a linear model from which we extract the slope in semi-log scale. We added more of these slopes to the set of summary statistics because it is known that the LTT slope contains information about the speciation and extinction rates for homogeneous diversification models (Nee et al. 1994; Ricklefs 2007). Lastly, for the BiSSE model we added the proportion of tips in state 1 (n_1_; and the proportion of tips in state 0 being simply n_0_ = 1 - n_1_).

| Name | Description | Number |
| --- | --- | --- |
| **Branch lengths** | | **25** |
| $\boldsymbol{height}$ | Height of the phylogeny: time from root to present | 1 |
| $<{bl}_{all}>$ | Mean length of all branches | 1 |
| $med({bl}_{all})$ | Median length of all branches | 1 |
| $var({bl}_{all})$ | Variance of the lengths of all branches | 1 |
| $<{bl}_{ext}>, med({bl}_{ext}), var({bl}_{ext})$ | Mean, median and variance of the lengths of external branches (*i.e.* linked to a tip) | 3 |
| $<{{bl}^{i}}_{int}>, med({{bl}^{i}}_{int}), var({{bl}^{i}}_{int})$ | Mean, median and variance of the lengths of internal branches (*i.e.* not linked to a tip) belonging to the $i\in\{1,2,3\}$ third of the phylogeny. | 9 (3x3) |
| $\frac{<{{bl}^{i}}_{int}>}{<{bl}_{ext}>}, \frac{med({{bl}^{i}}_{int})}{med({bl}_{ext})}, \frac{var({{bl}^{i}}_{int})}{var({bl}_{ext})}$ | Ratio of the piecewise mean, median and variance of internal branches respectively over the mean, median and variance of external branches. | 9 (3x3) |
| **Phylogeny topology** | | **8** |
| $\boldsymbol{colless}$ | Sum over internal nodes of the absolute difference between number of left and right tip(s) | 1 |
| $sackin$ | Sum over tips of the number of internal nodes separating the tip to the root. | 1 |
| $\frac{width}{depth}$ | Ratio of the maximal width over the maximal depth. Depth is defined as the number of branches separating a node from the root, and the width of a phylogeny at a given depth ($width(depth)$) is the number of nodes having that given depth. | 1 |
| $max(\Delta width)$ | Maximal width difference between two consecutive depths. | 1 |
| $max(ladder)$ | Maximal ladder size, where a ladder is a chain of internal nodes each linked to a single leaf. The ladder size is the number of internal nodes in the chain. | 1 |
| $\frac{internal node \in ladder}{number of internal nodes}$ | Proportion of internal nodes belonging to a ladder. | 1 |
| ${stairecaseness}_{1}$ | Proportion of internal nodes that have a different number of left and right tips (*imbalanced nodes*) | 1 |
| ${stairecaseness}_{2}$ | Mean of the ratio of the minimal number of tips on a side over the maximal number of tips on a side, for each internal node. | 1 |
| **LTT** | | **51** |
| $\boldsymbol{n}_{\boldsymbol{tips}}$ | Number of extant species | 1 |
| Binned LTT coordinates | LTT coordinates (time, number of lineages): binned the number of lineages in 20 equal parts and add the corresponding time. | 40 (20x2) |
| LTT slopes | Split the LTT in 10 equal parts regarding time and infer the slope from the linear model of the LTT (in semilog scale) for each part | 10 |
| **Additional statistic for BiSSE** | | **1** |
| $\frac{tips in state 1}{n_{tips}}$ | Ratio of number of tips in state 1 over the total number of tips | 1 |

**Table S0. List of summary statistics computed to summarize phylogenies.** These statistics are mainly inspired from (Saulnier et al., 2017).

### *Zero Padding of Phylogeny Representations*

Zero padding is a method used when one has data of different sizes and wants to have data of the same size. For instance, in our case the phylogeny representation sizes often depend on the size of the phylogenies. However, most neural network architectures need to be fed with data of the same size. Thus we need to use zero padding to overcome this issue. The principle of zero padding is to: 1) set a maximum data size; 2) for all the data with smaller sizes than the maximum size, fill with zeros until the data reaches the maximum size previously set. This process is detailed more precisely for the different neural network methods, as each method has its particularity.

#### *LTT with CNN.—*The LTT is a time series of N-1 points where N is the size of the phylogeny (or the number of tips). Therefore, since CNNs need inputs of the same sizes, we use zero padding to bring all LTT vectors to the same size of N_max_-1=999, where N_max_ is the maximum phylogeny size and N_max_ = 1,000.

*Phylogeny encoding with CNN.—*The phylogeny encoding has a size of 2N (N values for internal nodes plus phylogeny height + N values for the tips) for the CRBD model and 3N for the BiSSE model (+ N values for the tip states) with N the size of the phylogeny. For an arbitrary encoding vector of a phylogeny of size N, we do the following:

1) Extract the N first values corresponding to the N internal nodes plus height, and add zeros until having a vector of size N_max_;
2) Repeat for the N values corresponding to the tips;
3) If considering the BiSSE model, repeat for the N values corresponding to the tip states;
4) Concatenate the vectors resulting from the previous steps to obtain a final vector of size 2N_max_=2,000 for the CRBD model and 3N_max_=3,000 for the BiSSE model.

Lastly, when considering the BiSSE model the tip states are usually encoded using 0 and 1, however the use of 0 to encode for tip states can be confusing with the use of 0 for zero padding. Thus, we encode the tip states with -1 and 1 instead of 0 and 1 to avoid this ambiguity. We observed that encoding tip states as such led to a significant reduction of the total error.

*LTT with RNN.—*For RNN feeding on LTT, the zero padding was found to significantly deteriorate the RNN training, preventing the RNN from learning how to infer rates from LTTs. Thus we use a small trick to not use zero-padding for LTT with RNN. In the torch framework, RNNs can be fed with input data of different sizes, as long as in each batch data have the same sizes. Then for RNNs training, we created custom batches such that each batch contains LTTs of the same size. This trick might introduce a bias in the learning as the LTTs are not randomly given to the RNN, nevertheless, RNNs fed on LTT with this method have shown good inference performance and even outperform all other inference methods for the CRBD model.

Lastly, GNN with phylogeny graphs can work on input data of different sizes and summary statistics have always the same size (equal to the number of summary statistics considered), thus for DNN with summary statistics and GNN with phylogeny graphs zero padding is not needed.

### *Neural Network Structures and Parameters*

In this section, we describe the neural network structures (number of layers, number of hidden neurons) of the architectures studied and their main parameters (early stopping patience, dropout rate).

| **Layer name** | **Layer size** |
| --- | --- |
| Input | 86 (87) |
| Fully Connected 1 | 100 |
| Fully Connected 2 | 100 |
| Fully Connected 3 | 100 |
| Fully Connected 4 | 100 |
| Output | 2 |

**Table S1. Structure of the DNNs combined with summary statistics used to infer rate from diversification models.** In total, the network has 39,202 (39,302 if BiSSE) trainable parameters. In parentheses is the layer size if including tip state information in case of the BiSSE model.

| **Layer name** | **Layer size** |
| --- | --- |
| Input | 1,000 |
| Convolution 1 | 8*996 |
| Pooling 1 (average) | 8*498 |
| Convolution 2 | 16*494 |
| Pooling 2 (average) | 16*247 |
| Convolution 3 | 32*243 |
| Pooling 3 (average) | 32*121 |
| Fully Connected 1 | 3872 |
| Fully Connected 2 | 100 |
| Output | 2 |

**Table S2. Structure of the CNNs combined with LTTs used to infer rate from diversification models.** In total the network has 390,798 trainable parameters. The kernel size is equal to 5. For convolutional and pooling layers a size of 8*996 means that the layer has 8 channels and each channel contains a 1-d vector of length 996.

| **Layer name** | **Layer size** |
| --- | --- |
| Input | 2,000 (3,000) |
| Convolution 1 | 8*1,991 (8*2,991) |
| Pooling 1 (average) | 8*995 (8*1495) |
| Convolution 2 | 16*986 (16*1486) |
| Pooling 2 (average) | 16*493 (16*743) |
| Convolution 3 | 32*484 (32*734) |
| Pooling 3 (average) | 32*242 (32*367) |
| Convolution 4 | 64*233 (64*358) |
| Pooling 4 (average) | 64*116 (64*179) |
| Fully Connected 1 | 7,424 (11,456) |
| Fully Connected 2 | 100 |
| Output | 2 |

**Table S3. Structure of the CNNs combined with phylogeny encodings used to infer rate from diversification models.** In total, the network has 769,782 (1,172,982 if BiSSE) trainable parameters. In parentheses are the layer sizes if including tip state information in case of the BiSSE model. For convolutional and pooling layers a size of 8*996 means that the layer has 8 channels and each channel contains a 1-d vector of length 996.

| **Layer name** | **Layer size** |
| --- | --- |
| Input | 1,000 |
| LSTM 1 | 500 |
| LSTM 2 | 500 |
| Fully Connected | 500 |
| Output | 2 |

**Table S4. Structure of the RNNs combined with LTTs used to infer the rate from diversification models.** In total the network has 3,011,002 trainable parameters. The recurrent layers (LSTM 1 & 2) are composed of LSTM cells which is the most used type of cells for RNNs (for more details see Hochreiter and Schmidhuber, 1997). The recurrent layers had to be wide (500 neurons here) otherwise the inference was failing.

| **Layer name** | **Layer size** |
| --- | --- |
| Input | 5 (6) attributes per node |
| GraphConv 1 | 50 |
| GraphConv 2 | 50 |
| GraphConv 3 | 50 |
| GraphConv 4 | 100 |
| Fully Connected 1 | 100 |
| Fully Connected 2 | 50 |
| Output | 2 |

**Table S5. Structure of the GNNs combined with phylogeny graphs used to infer the rate from CRBD model.** The Graph Convolutional layer used was GCNConv (Kipf and Welling, 2017), other layer types (*e.g.* SAGEConv; see Hamilton et al., 2018) were tested but did not lead to any improvement. The GNN was implemented in Python with PyTorch Geometric (Fey and Lenssen 2019).

| **Layer name** | **Layer size** |
| --- | --- |
| Input | 5 (6) attributes per node |
| GraphConv 1 | 32 |
| GraphConv 2 | 32 |
| Fully Connected 1 | 32 |
| Output | 2 |

**Table S6. Structure of the GNNs combined with phylogeny graphs used to infer the rate from BiSSE model.** The Graph Convolutional layer used was GCNConv (Kipf and Welling, 2017), other layer types (*e.g.* SAGEConv; see Hamilton et al., 2018) were tested but did not lead to any improvement. The GNN was implemented in Julia with Flux (Innes 2018).

Moreover, to avoid overfitting of the neural networks, we used early stopping and dropout. For all architectures, we used a patience of 3 epochs for the early stopping, meaning that the neural network training stops when the total error on the validation has not improved since 3 epochs. Furthermore, we introduced a dropout rate p = 0.01, meaning that at each epoch 1% of all network weights are reset to 0.

### *Evaluating Deep-Learning Methods Performance Far From Boundaries of the Parameter Space*

Neural networks learn to not predict parameter values outside the parameter space it has seen during the training. Typically, if a neural network has only seen phylogenies with speciation rate in [0.1,1], the trained neural network will not predict speciation rate outside this range (see Fig. S1). This phenomenon artificially reduces the prediction error of the neural networks compared to the Maximum Likelihood Estimations that can go outside the simulated parameter space. Thus, to avoid this artifact, we restrict our analysis to an inner domain of the parameter space by excluding from the analysis the parameter values near the parameter space boundaries. The only exception to this is the lower boundary of extinction rate for the CRBD model, as in this case the Maximum likelihood is also constrained to only predict positive extinction rates. The parameter space as well as the inner domain considered to evaluate inference method performances are summarized in Table S6.

| **Diversification model** | **Rate** | **Parameter range** | **Inner range** |
| --- | --- | --- | --- |
| CRBD | Speciation rate | [0.1,1] | [0.2,0.9] |
|  | Extinction rate | [0,0.9] | [0,0.8] |
| BiSSE | Speciation rate | [0.1,1] | [0.2,0.9] |
|  | Transition rate | [0.01,0.1] | [0.02,0.09] |

**Table S7. Parameter space and inner domain considered for the evaluation of inference method performances.** The parameter range corresponds to the range in which the parameter was simulated. The inner range corresponds to the range which was selected to evaluate the method performances to avoid an artificial reduction of the prediction error for deep-learning methods.

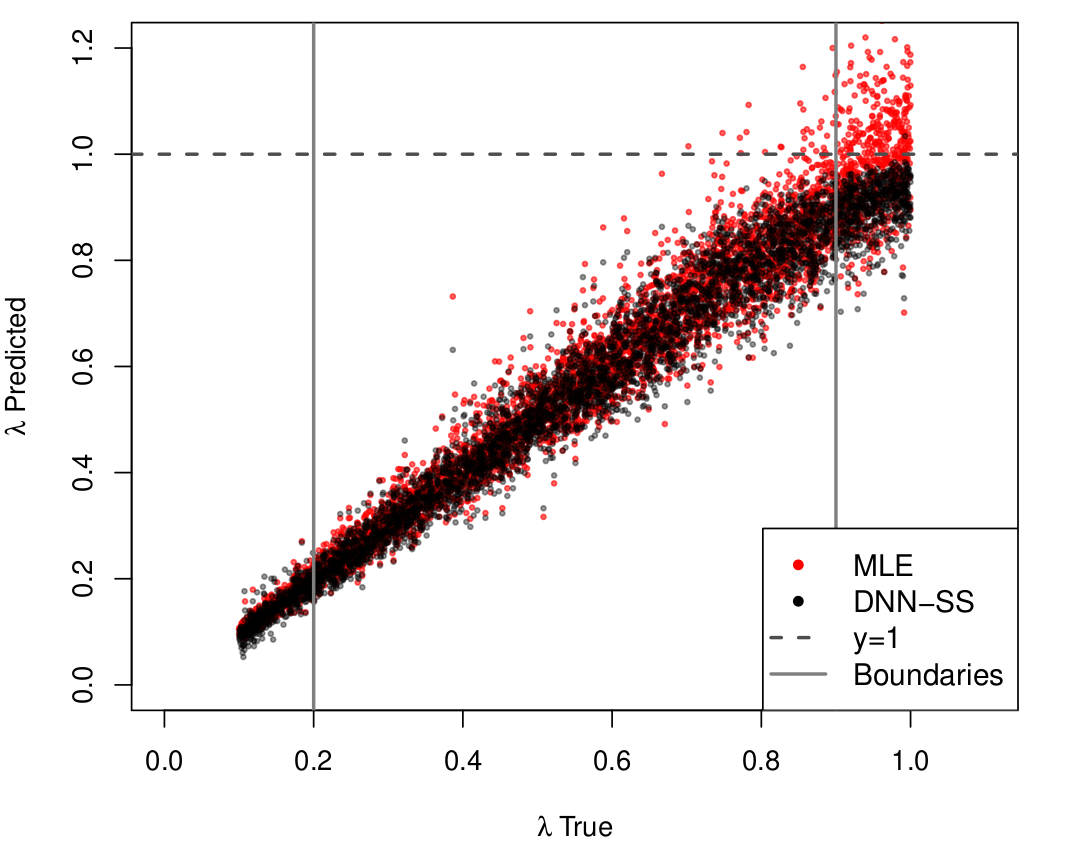

**Figure S1. Artificial reduction of the prediction error of deep learning methods due to parameter space boundaries.** Scatter plot of predicted vs. true values for the speciation rate ($\lambda$) of the CRBD model: predicted values by MLE are in red and predicted values by DNN fed with summary statistics (DNN-SS) in black. The true values of the speciation rate are uniformly distributed in [0.1,1]. We observe that the trained neural network learned to not predict speciation rate above 1 (horizontal dashed line) because it has learned the boundaries of the parameter space. This leads to an artificial reduction of the prediction error: ${SSD}_{DNN}= (6.2\pm0.3)\times{10}^{-3}$ if considering the whole range ($\lambda\in[0.1,1]$) while ${SSD}_{DNN}= (6.5\pm0.3)\times{10}^{-3}$ if considering an inner domain (between vertical plain lines; $\lambda\in[0.2,0.9]$).
